# Mutational signature profiling classifies subtypes of clinically different mismatch repair deficient tumors with a differential immunogenic response potential

**DOI:** 10.1101/2021.09.28.460630

**Authors:** Mar Giner-Calabuig, Seila De Leon, Julian Wang, Tara D Fehlmann, Chinedu Ukaegbu, Joanna Gibson, Miren Alustiza Fernandez, Maria-Dolores Pico, Cristina Alenda, Maite Herraiz, Marta Carrillo-Palau, Inmaculada Salces, Josep Reyes, Silvia P Ortega, Antònia Obrador, Michael Cecchini, Sapna Syngal, Elena Stoffel, Nathan A Ellis, Joann Sweasy, Rodrigo Jover, Xavier Llor, Rosa M Xicola

## Abstract

**Background:** Mismatch repair (MMR) deficiency is the hallmark of tumors from Lynch syndrome (LS), sporadic *MLH1* hypermethylated, and Lynch-like syndrome (LLS), but there is a lack of understanding of the variability in their mutational profiles based on clinical phenotypes. The aim of this study was to perform a molecular characterization to identify novel features that can impact tumor behavior and clinical management.

**Methods:** We tested 105 MMR-deficient colorectal cancer tumors (25 LS, 35 LLS, and 45 sporadic) for global exome microsatellite instability, cancer mutational signatures, mutational spectrum and neoepitope load.

**Results:** 78% of tumors showed high contribution of MMR-deficient mutational signatures, high level of global exome microsatellite instability, loss of MLH1/PMS2 protein expression and included sporadic tumors. 22% of tumors showed weaker features of MMR deficiency, 73% lost MSH2/MSH6 expression and included half of LS and LLS tumors. Remarkably, 9% of all tumors lacked global exome microsatellite instability. Lastly, HLA-B07:02 could be triggering the neoantigen presentation in tumors that show the strongest contribution of MMR-deficient tumors.

**Conclusions:** Next-generation sequencing approaches allow for a granular molecular characterization of MMR-deficient tumors, which can be essential to properly diagnose and treat patients with these tumors in the setting of personalized medicine.

## INTRODUCTION

The core of the mismatch repair (MMR) system includes two types of protein complexes that recognize nucleotide mismatches and insertion/deletion loops arising during DNA replication, MutS (MSH2/MSH6) and MutL (MLH1/PMS2), heterodimers(1). When the system is defective, DNA alterations occur throughout the genome. Microsatellites (short and repetitive sequences) are particularly prone to polymerase slippage and thus accumulate a significant number of mutations caused by MMR deficiency. In MMR-deficient tumors, the length of these sequence runs cannot be maintained during DNA replication, and cells show microsatellite instability (MSI)(2). MSI is commonly tested by amplification of five well-established mononucleotide markers. The identification of instability in two or more markers classifies a tumor as MSI-High (MSI-H), in one marker as MSI-Low (MSI-L) and in none as microsatellite stable (MSS)(2).

Three percent of all colorectal cancers (CRCs) develop in individuals with Lynch syndrome (LS)(3). LS is the most common inherited cancer syndrome and it is caused by germline pathogenic variants in an MMR gene (*MLH1, MSH2, MSH6, PMS2*)(4). In LS patients, cancer eventually develops when a second somatic mutation in the affected MMR gene is acquired resulting in biallelic MMR inactivation. About 15% of sporadic CRCs develop through biallelic somatic inactivation of *MLH1* following promoter hypermethylation(5). The *BRAF* gene mutation (V600E) is present in a third of these sporadic MSI tumors(6), but *BRAF* mutation is rarely seen in LS. Finally, a newer entity with MSI has recently been described as Lynch-like syndrome (LLS). The incidence of CRC is higher in families of LLS patients than in families with sporadic CRC, and their age of diagnosis is younger than in sporadic cases(7). The molecular complexity in LLS patients remains poorly understood. LLS patients’ MMR deficiency is neither due to germline mutations in MMR genes nor *MLH1* hypermethylation, and they do not have the V600E *BRAF* mutation. Previous studies showed that some of these LLS tumors have somatic mutations causing biallelic inactivation of an MMR gene(8-11).

Although the common final molecular characteristic of all three types of tumors is MMR deficiency, their mutational profiles can vary considerably. This has been observed through the study of mutational signatures, which are the unique combinations of mutation types that tumors accumulate due to several aberrant processes. Aberrant processes include infidelity of the DNA replication machinery, exogenous or endogenous mutagen exposures, enzymatic modification of DNA, or defective DNA repair. Mutational signature analysis can provide important insights into cancer development through the characterization of underlying mutational processes. To date, 77 mutational signatures have been described based on single base substitutions, doublet bases substitutions, and small insertion and deletions(12, 13). While the presence of single base substitution signatures SBS6, SBS14, SBS15, SBS20, SBS21, SBS26 and SBS44 (12, 13) has been associated with MMR-deficient tumors, no data explain how the variability of the presence of these signatures among these tumors can affect clinical features. Knowledge of these mutational differences could help refine the molecular definition of these tumors beyond MSI. This variability could have different implications on tumor behavior or treatment response. For example, the level of MSI and mutational burden can result in a different neoepitope load. Neoepitopes are the novo amino acid peptides generated from mutations that cancer cells acquire through the carcinogenic process. Cancer and immune cells present neoepitopes to T-cells promoting an immune response that eventually will kill the presenting cancer cell(14). Mutations that cause frameshift indels create a novel open reading frame and can produce a large number of neoepitopes highly distinct from self(15). Therefore, MSI tumors have high mutational burden associated with a significant neoepitope load and immune response. Leveraging the interactions among different immune response effectors is the molecular base for cancer immunotherapy. In that sense, cancer treatment of patients with MMR-deficient tumors has been revolutionized following demonstration of the effectiveness of immune checkpoint inhibitors such as pembrolizumab and ipilimumab/nivolumab in combination. The first clinical trial included 41 patients with a response rate of 40%(16). A second trial included 119 MMR-deficient CRC patients with a response rate of 54.6%(17). Thus, a significant number of patients with MSI tumors do not respond to these agents and tumor mutational burden has been suggested to more likely explain differences in response within this group(18). Still to be addressed is whether the presence of a specific mutational signature profile could improve the prediction of treatment response.

Because deeper molecular characterization of MMR-deficient tumors could better inform clinical management and targeted cancer treatment, we analyzed 105 MMR-deficient colorectal tumors. We compared global exome microsatellite instability, mutational spectrum, mutational signatures, and immunogenic response prediction between tumors that developed from sporadic *MLH1* hypermethylation/BRAF mutated tumors, LS and, LLS.

## MATERIALS and METHODS

### Study Population

We included 105 MMR-deficient tumors from 103 patients: 25 LS patients (4 *MLH1, 6 MSH2*, 7 *MSH6*, and 8 *PMS2* germline pathogenic variant carriers); 35 LLS patients; 43 patients with sporadic MSI/BRAF mutated tumors (Supplementary Table 1). We obtained DNA from two patients with multiple colorectal MSI tumors. DNA from sixty-five tumors was extracted from formalin-fixed paraffin-embedded tissue samples and 11 from fresh-frozen tissue samples (Supplementary Table 2). Patients were recruited through the EPICOLON Consortium(19), the Hospital General Universitario de Alicante in Spain, Yale New Haven Hospital, the Dana Farber Cancer Institute, the University of Michigan, and the Chicago Colorectal Cancer Consortium(20). Twenty-nine of the MSI/BRAF mutated tumors were obtained from the publicly available series of colon (COAD) and rectal (REAC) tumors from The Cancer Genome Atlas (TCGA) (RRID:SCR_003193). Clinical and demographic data for TCGA cases were obtained from their data repository(21). Each institution provided molecular, clinical information and germline genetic tests results. Patients were recruited in their original institutions under projects approved by institutional human research review committees. All data was shared in a de-identified manner to unlink any patient’s personal identifiers from their samples. Clinical characteristics and family history of cancer are summarized in Supplementary Table 1. A set of independent samples including six MMR-deficient tumors was collected including five LLS tumors and one MSI/BRAF mutated tumors. These samples were obtained from Institut d’Investigació Sanitària Illes Balears in Spain (Supplementary Table 3).

### Paired exome sequencing

Paired exome sequencing was performed at the Yale Center for Genome Analysis (YCGA). Reads were aligned to the GRCh37 (hg19) human reference, the first and last base from each read were trimmed. The Burroughs Wheeler aligner, BWA MEM, was used to align the paired-end reads. Samtools was used to sort the BAM file. Picard’s MarkDuplicates was used to mark PCR duplicates. The Genome Analysis Toolkit GATK 3 best practices workflow was used to realign indels, recalibrate base quality scores, generate GVCF files, call variants and filter variants using HardFiltering. Tumor and normal samples were computed separately, except for the variant calling step, which performed a joint calling with both tumor and normal samples. MuTect(22) computed the initial somatic point mutations. MuTect was given dbSNP v138 and COSMIC v72 to improve the identification of likely germline variants. ANNOVAR was run to include core annotations for each of the variants(23). In-house annotation identified variants in cancer genes from Oncomine, Foundation One or Cancer Gene Census gene lists. Non-coding and synonymous variants were filtered out unless they were within 15 bases of a splice site of a protein-coding gene. Any variant called as somatic was downgraded to germline based on minor allele frequency and the maximum population frequency for the variant. For each heterozygous variant called in the normal sample, where the depth of the normal sample is ≥ 20, the allele frequency difference was calculated between normal and tumor. A local polynomial regression was used to calculate the frequency difference data across the genome. Regions with fitted values > 0.1 were called as loss of heterozygosity (LOH) regions. The called LOH regions were manually curated to generate the final list of regions. Exome statistics for tumors are provided in Supplementary Table 4. Variants in cancer driver genes were only reported if they were predicted as nonsense, splicing and missense variants with a damaging effect score prediction in more than six of the ten algorithms used. For the TCGA set, we included nonsense, splicing and missense variants with a damaging effect score prediction in one of the two algorithms used. Before reporting any mutated genes, we excluded genes that are currently mutated in exome sequencing projects due to their size and the presence of repetitive sequences(24).

### MANTIS (Microsatellite Analysis for Normal Tumor InStability)

MANTIS(25) analyzes the instability of a normal-tumor sample pair as an aggregate of microsatellite loci instead of individual loci differences using next-generation sequencing data. The approach allows the tool to evaluate the global microsatellite instability present in a tumor sample, using the data from the corresponding normal sample as an error-correcting baseline. Furthermore, by pooling the scores of all the loci and treating the average as the instability score, the evaluation benefits from the law of large numbers, which reduces the impact that sequencing errors or poorly performing loci may have on the results. MANTIS showed higher performance in comparison to three other algorithms. MANTIS demonstrated the highest classification accuracy at a classification threshold of 0.4. We used the bam files from the paired normal and tumor samples. We used the parameters --threads 4 -mrq 20.0 -mlq 25.0 -mlc 20 -mrr 1, as recommended per exome sequencing usage. MANTIS score Step-Wise ≥ 0.4 was considered unstable. Tumors with a MANTIS score < 0.4 were classified as global microsatellite stable.

As the vcf files for paired samples from the TCGA were not available to us, we used the MANTIS score classification published by Bonneville R et al.(26) for the 29 TCGA MSI/BRAF mutated samples included in our study.

### MSIseq: Assessing Microsatellite Instability from Catalogs of Somatic Mutations

MSIseq(27) software classifies tumors based on their MSI status. MSIseq uses the list of somatic single-nucleotide substitutions and microindels mutations generated from next-generation sequencing of whole exomes to classify tumors based on their MSI status. To develop the MSI-classifier in the software, the authors used four machine learning frameworks (logistic regression, decision tree, random forest, and naïve Bayes) to analyze a set of variables, including the total length of the sequence that was targeted for hybridization capture, information on mutations, and the type of cancer. Among the frameworks, the decision tree exhibited the highest concordance with laboratory tests. From all the variables, the final MSI-classifier takes into account the number of microindels in simple sequence repeats per Mb, named S.indv. Tumors with S.ind_D_>_D_0.395 are classified as MSIH. We used this approach as an alternative methodology to classify tumors for their MSI status. MSIseq differentiates from MANTIS because MSIseq uses a list of somatic mutations but MANTIS uses bam files for the normal and tumors samples. MSIseq is available as an open-source R package. We classified tumors in our cohort following the procedure described in the package’s vignette using the classifier provided by the package and the cancer type of our samples.

### Identification of known mutational signatures

We used the R package “MutationalPatterns”(28) to create the 96-trinucleotide mutation count matrix based on single base substitutions from each somatic vcf file. We then used a function (“fit_to_signatures”) that finds the optimal linear combination of mutational signatures that most closely reconstructs the mutation matrix by solving a non-negative least-squares constraints problem. The result is a matrix of the contribution from each of the 60 SBS COSMIC signatures (Version 3.2-March 2021) per sample. We followed the analysis steps described in the package’s vignette. The normalized contribution matrix was used to perform unsupervised hierarchical clustering using the Euclidean distance and a complete linkage algorithm (R packages: ComplexHeatmap(29) and dendsort(30)).

### Neoepitope prediction using NeoPredPipe

NeoPredPipe is an open source package (31) that allows users to process neoantigens predicted from vcf files using ANNOVAR(23) (RRID:SCR_012821) and netMHCpan(32) (RRID:SCR_018182). The first step of the analysis is to use an external software, POLYSOLVER(33), to genotyping HLA based on whole exome sequencing data and infers alleles for the three major MHC class I (HLA-A, -B, -C) genes. NeoPredPipe starts the pipeline using ANNOVAR to predict the mutated amino acid sequence from annotated nonsynonymous variant calls and the peptide sequence surrounding the newly introduced amino acid is extracted for epitope prediction. Then, the user can specify the epitope lengths to conduct predictions for typical epitopes of 8-, 9-, or 10-mers, using netMHCpan software. Based on patient’s HLA alleles and the somatic mutations identified, netMHCpan delivers candidate information for neoantigens from provided variant calls that may be presented to T-cells efficiently with strong affinity. NeoPredPipe incorporates the use of PeptideMatch(34) to check predicted neoepitopes for novelty against the human reference proteome. Afterward, NeoPredPipe employs the algorithms and process utilized by Luksza M et. al. (35) to calculate the likelihood of a neoantigen eliciting an immune response, thus providing a recognition potential estimate.

### Statistical Comparisons

For comparison between categorical variables, we used Fisher’s Exact test. For comparison of continuous variables, we used the Wilcoxon non-parametric test. All statistical comparisons and graphical representations were performed using R for statistical computing(36). P-values were adjusted for multiple comparisons using the Benjamini & Hochberg method.

## RESULTS

### Three distinct groups of tumors were identified based on the contribution of MMR-deficient mutational signatures

Through hierarchical clustering we identified groups or clusters of tumors that share a similar contribution of mutational signatures. We noticed that SBS87 was driving the clustering of the samples accordingly to the molecular events generated by thiopurine treatment (Supplementary Figure 1). As the aim study was to unveil the contribution of signatures underlying the carcinogenic process but not as the result of treatment, we removed this signature from the analysis.

Three main clusters emerged based on the level of contribution from 60 known Single Base Substitutions (SBS) cancer mutational signatures. Cluster 1 included 65 tumors (62%), Cluster 2 included 23 tumors (22%) and Cluster 3 the remaining 17 tumors (16%) (Figure 1). The main signatures identified correlated with well-known signatures related to MMR-deficient tumors, signature SBS6, SBS15, SBS20, SBS26 and the unspecific signature SBS1. We provide the mutational signature profile in a bar plot for each sample, the mutational profile for each sample, and the contribution of indels and single nucleotide variants as supplementary material (Supplementary Figure 2, 3 & 4).

**Figure 1.**
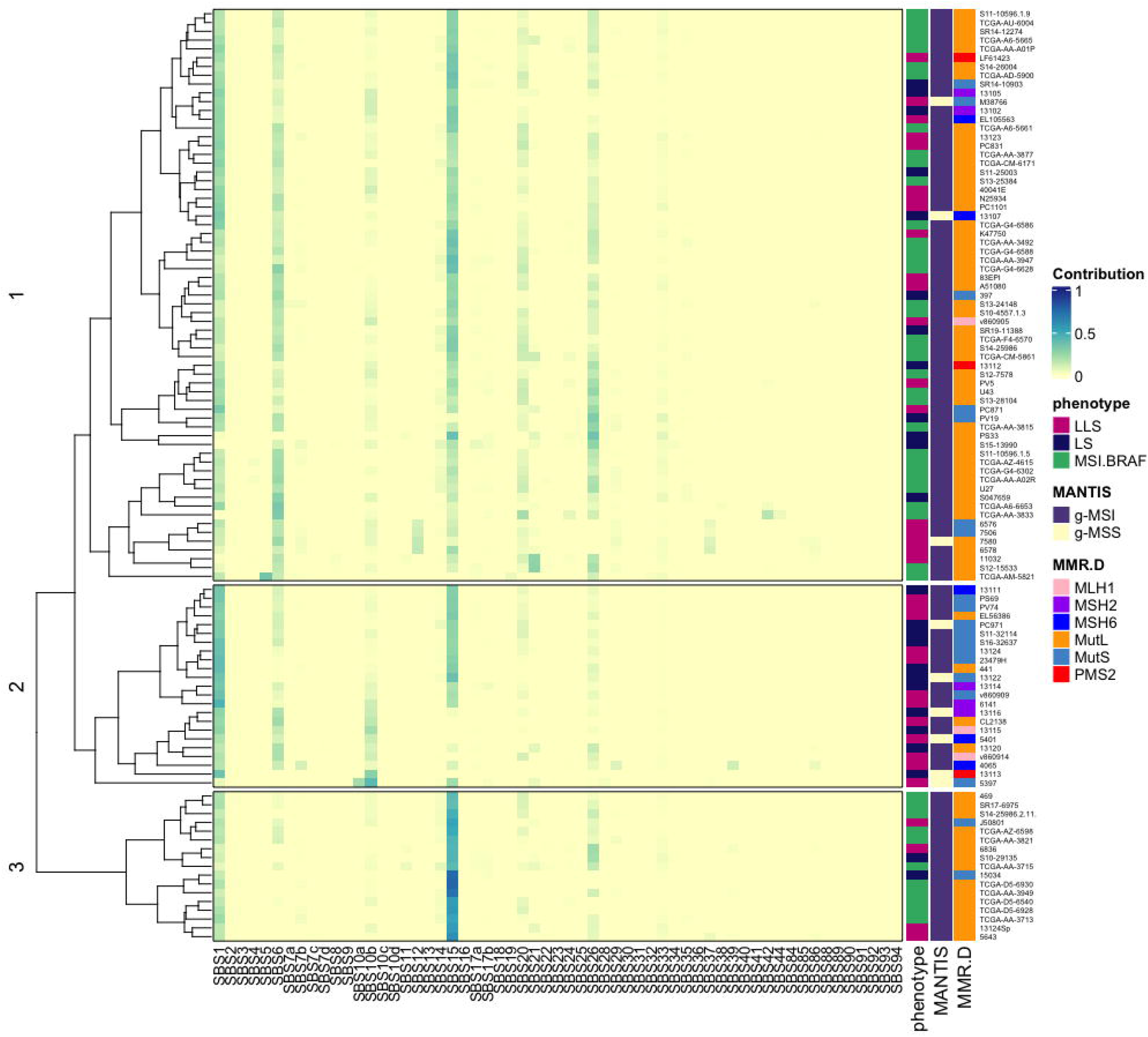
Mutational signature profile of MMR-deficient tumors. Clustering representation of the contribution from mutational signatures to each tumor. Each row represents one tumor and each column one signature. Annotation describes phenotype, loss of MMR protein complex expression, MANTIS classification. SBS87 is the result of thiopurine treatment. This signature was contributing to the mutational profile of a significant number of samples. As the aim study was to unveil the contribution of signatures underlying the carcinogenic process and not treatment, we remove it from the analysis. LLS = Lynch-like syndrome, LS = Lynch syndrome, MSI.BRAF = Microsatellite unstable with BRAF V600E, g-MSS= global microsatellite stable MANTIS<0.4, g-MSI= global microsatellite unstable MANTIS>0.4, MMR.D = loss of MMR protein expression, MutL = loss of expression of MLH1 and/or PMS2, MutS = loss of expression of MSH2 and/or MSH6.

In comparison to Cluster 2, Cluster 1 and 3 exhibited a significantly higher contribution of MMR deficient-related signatures, showed a significantly higher number of frameshifts mutations, and had a higher MANTIS score (Table 1). These features suggested that tumors in Cluster 1 and 3 had a significantly higher level of global exome microsatellite instability and MMR-deficient features. Cluster 1 was characterized by tumors with an equally high contribution from signatures SBS1, SBS6, and SBS15. Cluster 3 had the highest contribution of SBS15 but no contribution of SBS6. Cluster 1 and 3 were significantly enriched (83%) with tumors that lost the expression of the MutL complex (Table 1).

**Table 1.** Comparison of clinical and molecular features of tumors between clusters. LLS = Lynch-like syndrome, LS = Lynch syndrome, MSI.BRAF = Microsatellite unstable with BRAF V600E, VUS = variant of uncertain significance, MutL = loss of expression of MLH1 and/or PMS2, MutS = loss of expression of MSH2 and/or MSH6, M = male, F = female, CRC = colorectal, L = CRC left-sided, R = CRC right-sided, FH = family history, ≥ 1FDR = One or more first degree relative, TMB=Tumor Mutational Burden (total mutations/Mb); Sig = signature, ns = *FDR* >.05.

|  |  | Cluster1 | Cluster2 | Cluster3 | FDR |
| --- | --- | --- | --- | --- | --- |
|  | Patients | n=64 | n=23 | n=16 |  |
| **Phenotype** | LLS | 30% (19) | 48% (12) | 25% (4) | 3.62x10<sup>-5</sup> |
|  | LS | 19% (12) | 48% (11) | 13% (2) |  |
|  | MSI.BRAF | 51% (33) | 0% (0) | 63% (10) |  |
| **Sex** | M | 42% (27) | 52% (12) | 50% (8) | ns |
|  | F | 58% (37) | 48% (11) | 50% (8) |  |
| **Age**<br>**(median ±sd)** |  | 68±16 | 50±13 | 77±8 | 0.0028 |
| **Location** | L | 17% (11) | 30% (7) | 6% (1) | 0.0155 |
|  | R | 78% (50) | 48% (11) | 94% (15) |  |
|  | NA | 4% (3) | 22% (5) | 0% (0) |  |
| **Stage** | I/II | 61% (39) | 43% (10) | 81% (13) | 0.0098 |
|  | III/IV | 25% (16) | 17% (4) | 19% (4) |  |
|  | NA | 14% (9) | 39% (9) | 0% (0) |  |
| **FH of CRC** | none | 69% (40) | 58% (11) | 86% (12) | ns |
|  | >1FDR | 31% (18) | 42% (8) | 14% (2) |  |
|  | Tumors | n=65 | n=23 | n=17 |  |
| **Loss Expression MMR\*** | MutL | 83% (53) | 27% (6) | 88% (14) | 2.91x10<sup>-5</sup> |
|  | MutS | 17% (11) | 73% (16) | 12% (2) |  |
| **Signature**<br>**Composition (%)** | SBS1 | 236±187 | 507 ±315 | 199 ±73 | 0.00081 |
|  | SBS6 | 200±132 | 126±111 | 0±54 | 2.91x10<sup>-5</sup> |
|  | SBS14 | 15±29 | 10±37 | 33±41 | ns |
|  | SBS15 | 289±262 | 414±412 | 751±292 | 3.62x10<sup>-5</sup> |
|  | SBS20 | 81±62 | 37±77 | 15±88 | 0.0113 |
|  | SBS21 | 0±156 | 0±82 | 0±33 | ns |
|  | SBS26 | 113±152 | 49±45 | 69±106 | 0.0045 |
|  | SBS44 | 0±2 | 0±0 | 0±0 | ns |
| **Tumor purity (%)** |  | 55±13 | 50±16 | 60±14 | ns |
| **# events**<br>**(median ±sd)** | TMB | 40± 20 | 45±38 | 43±21 | ns |
|  | frameshifts | 197±133 | 68±100 | 219±223 | 0.0039 |
| **Mantis score** |  | 0.66±0.26 | 0.48±0.16 | 0.69±0.25 | 0.0028 |

\*one tumor in Cluster 2 did not show loss of MMR expression, but was MSI-H

Tumors in Cluster 2 showed a weaker level of MMR-deficient features. These tumors showed a significantly higher contribution from signature SBS1 than tumors in Cluster 1 and 3. Although tumors in Cluster 2 have a relatively high contribution of SBS15, the overall contribution of MMR-deficient signatures to the mutational profile was lower than in Cluster 1 and 3. Additionally, 73% of tumors in Cluster 2 lost expression of the MutS complex.

As tumors in Clusters 1 and 3 showed a significantly stronger level of microsatellite instability and the presence of frameshift mutations could be influenced by the level of tumor cells present in the DNA sample analyzed, we compared the tumor purity among tumors in each cluster. We did not identify a significant difference in the median tumor purity between tumors in each cluster (54 for Cluster 1, 50 for Cluster 2 and 60 for Cluster 3) (Table 1).

### Clinical differences between patients in each cluster

All tumors with the V600E *BRAF* were in either Cluster 1 or 3, representing 51% of tumors in Cluster 1 and 63% in Cluster 3. Thirty percent of tumors in Cluster 1 were LLS tumors and 19% were LS. In Cluster 3, 25% of tumors were LLS and 13% were LS whereas in Cluster 2 tumors were equally split between LS and LLS. Tumors in Cluster 1 and 3 were located predominantly on the right-side of the colon. Patients with these tumors were diagnosed significantly later than tumors in Cluster 2 with a median age of 68 and 77 in comparison to 50, respectively (Table 1). These differences were driven by the enrichment of patients with V600E *BRAF* mutated tumors in Cluster 1 and 3. Patients with tumors with the V600E *BRAF* mutation were diagnosed with a median age of 79 years old and 98% were located on the right-side of the colon (Supplementary Table 1). Excluding patients with *BRAF* V600E tumors from Cluster 1 and 3, we did not identify clinical differences between clusters (Supplementary Table 5).

### Biallelic inactivation of MMR genes

Loss of function of an MMR gene results from inactivating defects affecting both alleles. Therefore, all tumors with loss of protein expression of an MMR gene might have an identifiable biallelic inactivation. It is believed that *MLH1* promoter methylation is a biallelic event, and that LS tumors develop after acquiring a somatic mutation inactivating the second allele. However, the frequency of double somatic inactivation in LLS tumors is still not well-established. In our series, the frequency of double somatic inactivation for LLS was 54%, leaving about half of the LLS cases without obvious molecular biallelic inactivation triggering MMR deficiency. Among tumors lacking a biallelic inactivation, 48% had somatic mutations in different MMR genes, 30% had only one mutation and 22% lacked mutations in an MMR gene. The frequency of biallelic inactivation among LS tumors was 88%. Only three tumors from LS patients did not have an acquired somatic defect. Interestingly, all three tumors were from patients harboring a germline mutation in *PMS2*.

### Tumors with global exome microsatellite stability

Out of 105 tumors, nine were classified as global exome microsatellite stable based on their MANTIS score <0.4. Six of the nine tumors were in Cluster 2 and three in Cluster 1. To provide an alternative measure of microsatellite instability we used MSIseq to classify these nine tumors again. All the tumors were classified as non-MSI by this software, corroborating the MANTIS classification. Four of the tumors were LLS and five were LS tumors. Only two tumors lost expression of MutL and four of the nine tumors lost only MSH6 expression. Only one tumor would not be classified as hypermutated based on tumor mutation burden <10 mut/Mb (Supplementary Table 6).

Interestingly, one of the global exome microsatellite stable tumors with loss of expression of MSH6 (ID:5397) showed a clear different signature contribution in Cluster 2. This tumor separated from the rest through a single ramification, and it showed 60% contribution of signature 10 (20% SBS10a and 40% SBS10b). This tumor had the highest tumor mutational burden (193 mut/Mb). Signature 10 has been associated with mutations in the *POLE* gene. This tumor had a well-known hotspot mutation in the exonuclease domain of *POLE* (p.V411L) associated with an MSS hypermutator phenotype, corroborating our classification.

We sought to include an independent series of MMR-deficient tumors to validate our findings. We analyzed five LLS tumors and 1 MSI/BRAF mutated tumor (Supplementary Table 3). One LLS tumor was classified as global exome microsatellite stable, and it clustered together with the other global exome microsatellite stable tumor with loss of PMS2 from the original series. Another tumor from the original series with loss of PMS2 expression also cluster together with these two tumors in Cluster 2. The remaining five tumors, including the MSI/BRAF tumor, were in Cluster 1 and were global exome microsatellite unstable (Supplementary Figure 4).

### Differences in Cancer Driver mutations between tumors in each cluster

We further investigated mutational differences between these groups of tumors. We identified somatic mutations and LOH events in colorectal cancer driver genes (37). We included loss-of-function mutations or missense mutations predicted damaging by 6 or more of the 10 bioinformatics algorithms used. Tumors in Cluster 1 showed a higher frequency of somatic events in all cancer genes analyzed except for *AMER1* and *ARID1A*, which their mutation frequency was higher in Cluster 3. Differences did not reach statistical significance (Supplementary Table 7). Tumors from the TCGA series were excluded because data for LOH was not available.

### Differences in neoepitope presentation between tumors in each cluster

Our results described three types of tumors with different levels of microsatellite instability resulting in differences in the number of frameshift mutations and a different mutational spectrum and contribution from MMR-deficient mutational signatures. These molecular differences could affect the neoepitope load and, therefore, result in a different immunologic response. We performed a neoepitope prediction analysis using each patients’ HLA allele genotypes and the somatic mutations identified in 73 samples to test this hypothesis. We compared the most common alleles for each HLA-I gene and we did not identify any significant difference among the patients in each cluster (Supplementary Table 8). However, we identified that some of the alleles had nearly the same frequency for each genotype, suggesting linkage disequilibrium. To verify that, we used the database from the National Marrow Donor Program (https://bioinformatics.bethematchclinical.org/hla-resources/haplotype-frequencies/high-resolution-hla-alleles-and-haplotypes-in-the-us-population/) that provides the frequency of the most common HLA-I haplotypes among different ethnic populations. B07:02-C07:02 is present in 13% of White-European individuals. In our series, this haplotype represented 23% of individuals. B07:02-A02:01 is present in 5% of White-European individuals and in our series represented 11% of individuals. Based on the information reported by the patients included in the study, no ethnic differences were identified between clusters and 80% of individuals in each cluster were White-European.

#### Neoantigen prediction from frameshift mutations

We identified the total number of novel neoantigens arising from frameshift mutations that bind with strong affinity to a specific HLA-I with a high potential (likelihood) of inducing an immune response through a T cell receptor activation. We ranked all the interactions based on the likelihood score and identified the ones at the top 10% and the ones at the lower 10%. To approach this analysis, our working hypothesis was that if there was no selection towards a specific HLA -I allele, the distribution of alleles among different levels of likelihood should not be significantly different than the distribution of HLA-I alleles among the population of patients in each cluster. To test that hypothesis, we compared the distribution of HLA-I alleles between the population of patients in each cluster and the distribution of HLA-I alleles contributing to in each level of likelihood. Among the top 10%, B7:02 and A02:01 represented more than half of the interactions (42% and 21%, respectively). When we stratified by cluster and compared the allele distribution to the patient population allele frequency, we identified that Cluster 1 and 3 were significantly enriched with B7:02-neoantigens compared to Cluster 2 (Table 2). A02:01 did not show a significant difference in the patient population frequency. As we mentioned, B7:02 can be linked to C07:02 among White-Europeans. Interestingly, C07:02 was not contributing to any of the high potential HLA-neoantigens interactions. Among interactions in the lowest 10%, C07:02 contributed the most with 14%, B7:02 contributed 11%, and A02:01 contributed 3%. For the lowest 10%, the contribution of A02:01 was significantly depleted. These data suggest that B7:02 can trigger the neoantigen presentation in tumors that show the strongest contribution of MMR-deficient features (Clusters 1 and 3). It could be modulating a strong induction of immune response.

**Table 2:** HLA-I alleles – neoepitope interactions

| Cluster | B07:02 allele population | B07:02- frameshift neoepitopes interactions |  |  |  | B07:02- missense neoepitopes interactions |  |  |  |
| --- | --- | --- | --- | --- | --- | --- | --- | --- | --- |
|  |  | Top 10% | FDR | Low 10% | FDR | Top 10% | FDR | Low 10% | FDR |
| 1 | 14/74 (16%) | 88/104 (46%) | $6 \times 10^{-6}$ | 21/174 (11%) | ns | 32/142 (18%) | ns | 10/146 (6.4%) | ns |
| 2 | 3/41 (7%) | 15/51 (23%) | ns | 6/49 (11%) | ns | 4/118 (3%) | ns | 1/99 (1%) | ns |
| 3 | 3/15 (17%) | 39/42 (48%) | 0.044 | 9/63 (12%) | ns | 6/57 (9.5%) | ns | 3/59 (4.8%) | ns |

| Cluster | A02:01 allele population | A02:01- frameshift neoepitopes interactions |  |  |  | A02:01- missense neoepitopes interactions |  |  |  |
| --- | --- | --- | --- | --- | --- | --- | --- | --- | --- |
|  |  | Top 10% | FDR | Low 10% | FDR | Top 10% | FDR | Low 10% | FDR |
| 1 | 17/71 (19%) | 40/152 (21%) | ns | 3/192 (1.5%) | $3.2 \times 10^{-6}$ | 42/132 (24%) | ns | 0/156 (0%) | $1.8 \times 10^{-8}$ |
| 2 | 12/32 (27%) | 9/57 (14%) | ns | 4/51 (7.3%) | 0.033 | 29/93 (23.7) | ns | 1/99 (1%) | $1.8 \times 10^{-5}$ |
| 3 | 7/11 (39%) | 24/57 (30%) | ns | 3/69 (4.2%) | 0.0016 | 28/35 (44%) | ns | 0/64 (0%) | $3.6 \times 10^{-8}$ |

| Cluster | C07:02 allele population | C07:02- frameshift neoepitopes interactions |  |  |  | C07:02- missense neoepitopes interactions |  |  |  |
| --- | --- | --- | --- | --- | --- | --- | --- | --- | --- |
|  |  | Top 10% | FDR | Low 10% | FDR | Top 10% | FDR | Low 10% | FDR |
| 1 | 14/74 (16%) | 0/192 (0%) | $7.6 \times 10^{-7}$ | 32/163 (16%) | ns | 2/172 (1.1%) | $3.6 \times 10^{-5}$ | 11/145 (7%) | ns |
| 2 | 2/42 (45%) | 0/66 (0%) | ns | 1/54 (1.8%) | ns | 0/122 (0%) | ns | 1/99 (1%) | ns |
| 3 | 4/14 (22%) | 0/81 (0%) | 0.0029 | 12/60 (17%) | ns | 0/63 (0%) | 0.00648 | 1/63 (1.6%) | 0.02247 |

#### Neoantigen prediction from missense mutations

We performed the same analysis, including the neoantigens resulting from missense somatic mutations. Similar results were observed, but B7:02 allele was not significantly driving the interactions among the top 10% (Table 2).

## DISCUSSION

This study aimed at describing the heterogeneity of MMR-deficient tumors and establishing unique molecular features and clinical correlates.

We found that MMR-deficient tumors with MSI, as defined by conventional MSI PCR and/or IHC of MMR proteins, are a heterogeneous group. Using next-generation sequencing data we identified three types of MMR-deficient tumors based on the level of MSI. While technical limitations could have affected the sequencing and identification of somatic variants, tumors included in the study had a median of 105 mean coverage and we did not include any tumor with tumor cell content lower than 20% (Supplementary Table 4). Discrepancies could also result from mutational heterogeneity of the different areas in the tumor(38). Nevertheless, the use of next-generation sequencing can provide a broader determination of the real status of microsatellite instability, beyond the common analysis of just five microsatellites as done by traditional PCR. We hypothesize that among MSI tumors there is a group of tumors that show loss of MMR protein expression, but still retain MMR functionality and thus develop some degree of microsatellite instability but not global microsatellite instability. We believe that the classification of 10 tumors as globally exome microsatellite stable is accurate because these tumors have a very low number of frameshift mutations, but a higher than the average number of mutations in tumors classified as MSS by standard PCR(39). Moreover, one tumor was already identified as MSI-L by PCR. Therefore, next-generation sequencing would be a suitable technique for identifying these not strictly microsatellite unstable tumors. Further research is required to clarify these cases and the implication that this type of tumors might have in tumors with loss of expression of the complex MutS in the setting of LS and LLS tumors. In fact, Latham A. et al(40) described that approximately 36% of LS patients had an exome globally microsatellite stable phenotype.

With the current study, we correlated the level of instability and mutational signatures with clinical MMR phenotypes. With doing so, we were able to replicate several aspects that in-vitro studies have shown in knockout cell lines (41, 42). Zou et. al. demonstrated that *PMS2* knockout cell lines had a differential mutational signature (42). We observed that three out of the four patients with mutations and loss of expression only in this protein clustered together demonstrating a similar signature contribution. Additionally, Zou et. al. showed that *MSH6* knockout cell line had a high contribution of single nucleotide variants but fewer indels. The effect of this different mutational profile can be contributing to the global exome microsatellite stable subgroup of tumors, as four out of nine had only mutations in *MSH6* and loss of protein expression. In future studies it will be essential to frame the existence of these specific molecular types in the overall screening for LS patients.

Another novel aspect of this study was the analysis of LLS tumors. As with LS tumors, we showed that this group of tumors is also heterogeneous according to their level of instability. We report that in 46% of LLS patients we were not able to identify a somatic biallelic inactivation of one of the MMR genes. In comparison, 12% of the LS cases did show biallelic inactivation. This means that nearly half of the tumors from LLS patients do not have an evident molecular explanation for the presence of MSI, suggesting that other poorly understood mechanisms might contribute to the development of MSI in these cases. A potential explanation could be an epigenetic event. *Buckley et al*. (43) described a novel association between methylation of *SHPRH* gene and MSI burden and they also reported that 24% of MMR-deficient tumors lacked MMR mutations, which is similar to the frequency we identified among LLS tumors. Unfortunately, in this series we were not able to evaluate the role of promoter methylation and gene transcription silencing in the molecular heterogeneity of the tumors in our study. We(20) and other authors(11, 44) have described germline mutations in other DNA repair genes as being associated with the LLS phenotype, which could contribute to increased genomic instability, mutational load and subsequent mutations in the MMR genes.

Nevertheless, our findings’ most relevant clinical implication is that mutational signatures identify tumors with a higher level of microsatellite instability, which correlates with a potentially stronger immunogenic response. The observed difference in potential immunogenic response could be influenced by the mutational load and also differences in HLA-I alleles between patients. Our results are highly relevant for the treatment of patients with MMR-deficient tumors. Even though MMR-deficient tumors have a high mutational burden and should respond to immune checkpoint inhibitors(45), not all of them do. In fact, the intensity of instability identified in MMR-deficient tumors has been suggested to impact the response to immune checkpoint inhibitors(46) and explains, in part, the inconsistent response to immunotherapy. Moreover, recent data also show how HLA-I alleles determine the degree of response to immunotherapy. In a study of patients with melanoma tumors, the authors showed how maximal heterozygosity at HLA-I loci improved overall survival after immunotherapy treatment. HLA-I homozygosity and low mutation burden were strongly associated with decreased survival compared with patients who were heterozygous at each class I locus and whose tumors had high mutation burden(47). Lastly, our current study suggests that specific HLA-I alleles, i.e HLA-B7:02, could promote stronger immunogenic response. This could be explained by differences in specific structural features that may modulate the effective T cell recognition of neoepitopes presented by this allele(47).

Genetic differences among populations could influence variation in HLA-I genotypes, but we do not believe that this is the case. In the current study, 80% of individuals in each cluster were White-European. The mentioned haplotype, B07:02-C07:02, is less frequent among African Americans (49), representing our series’s second most common ethnicity.

Immune checkpoint inhibitors are approved for the refractory treatment of MMR-deficient or MSI tumors. Patients with these tumors are currently being treated with immunotherapy regardless of tumor type or additional molecular features. Our study demonstrates the value of whole exome sequencing as a unique test to properly characterize CRC tumors in terms of global exome microsatellite instability, mutational signatures, neoepitope load and HLA-I genotyping, which may identify patients who would benefit from immunotherapy treatment, more accurately. Exome sequencing of matched normal and tumor DNA would also diagnose patients for LS, identify LLS patients and identify tumors with double somatic mutations in an MMR gene.

Our study has the obvious limitation that patients in our study were not treated with immunotherapy. Our patient cohort is retrospective and immunotherapy treatment has been only recently implemented in the clinic. Future studies will include MMR-deficient CRC patients treated with immune checkpoint inhibitors to perform a direct comparison of treatment response among molecular subgroups. Finally, we believe that our study provides relevant information on how the determination of the molecular heterogeneity among MMR-deficient CRC tumors is essential to properly diagnose and treat these patients in the setting of personalized medicine.

## Acknowledgements

The results shown here are in part based on data generated by the TCGA Research Network: https://www.cancer.gov/tcga.

## Author Contributions

Conceptualization (MGC,XL,RMX); Data curation (MGC,SDL,JW,MDP,TF,CU,MH,MCP,IS); Formal analysis (MGC,RMX); Funding acquisition (RJ,JS,XL,RMX); Investigation (MGC,SDL,JW,JG,CA); Project administration (RMX); Resources (TF,CU,SS,MH,MCP,IS,EMS,NAE,JS,MA,MDP,RJ,JR,SPO,AO); Supervision (RMX); Validation (MGC,SDL); Visualization (MGC,JW,RMX); Writing – original draft (MGC,XL,RMX); Writing – review & editing (MGC,CU,SS,ES,JG,MC, RJ,NAE,XL,RMX).

## Ethics approval and consent to participate

Patients were recruited and consented in their original institutions under projects approved by institutional human research review committees. At Yale University, the project was approved by the Biomedical Institutional Review Board. All data was shared in a de-identified manner to unlink any patient’s personal identifiers from their samples. The study was performed in accordance with the Declaration of Helsinki.

## Data availability

The datasets generated and/or analyzed during the current study are not publicly available yet but are available from the corresponding author on reasonable request.

## Competing interests

Dr. Syngal is a consultant for Myriad Genetics and DC Health Technologies and has rights to an inventor portion of the licensing revenue from PREMM5. The rest of the authors have nothing to disclose.

## Funding information

This work was supported by grants from the National Cancer Institute (1K01CA204431-01A1 R.M.X), the Prevent Cancer Foundation (RMX), Colorectal Cancer Alliance’s Chris4Life grant (RMX), pre-doctoral grant from Conselleria d’Educació de la Generalitat Valenciana. VALi+d. EXP ACIF/2010/018, ACIF/2016/002 (MGC), Instituto de Salud Carlos III PI17/01756 (RJ), Asociación Española de Gastroenterología. Beca Tamarite 2017 (RJ) and the Donaldson Foundation (SS and CU).

## Notes

### Competing Interest Statement

The authors have declared no competing interest.

