## Supplementary material for "Mutational signature profiling classifies subtypes of clinically different mismatch repair deficient tumors with a differential immunogenic response potential"

**Supplementary Figure 1. Mutational signature profile of MMR-deficient tumors including all signatures**

Clustering representation of the contribution from mutational signatures (COSMIC v3.2) to each tumor. Each row represents one tumor and each column one signature. Annotation describes phenotype, loss of MMR protein complex expression, MANTIS classification.

LLS = Lynch-like syndrome, LS = Lynch syndrome, MSI.BRAF = Microsatellite unstable with BRAF V600E, g-MSS= global microsatellite stable MANTIS<0.4, g-MSI= global microsatellite unstable MANTIS>0.4, MMR.D = loss of MMR protein expression, MutL = loss of expression of MLH1 and/or PMS2, MutS = loss of expression of MSH2 and/or MSH6.

**Supplementary Figure 2. Mutational signature contribution represented in a barplot for each sample**

**Supplementary Figure 3. Representation of the 96 bar plot mutational profile for each tumor**

**Supplementary Figure 4. Insertion/Deletion (InDels) and single nucleotide variants (SNVs) contribution to the tumor mutational burden for each sample by cluster**

**Supplementary Figure 5. Mutational signature profile including the independent set of dMMR tumors.**

Clustering representation of the contribution from mutational signatures (COSMIC v3.2) to each tumor. Each row represents one tumor and each column one signature. Annotation describes phenotype, loss of MMR protein complex expression, MANTIS classification.

LLS = Lynch-like syndrome, LS = Lynch syndrome, MSI.BRAF = Microsatellite unstable with BRAF V600E, g-MSS= global microsatellite stable MANTIS<0.4, g-MSI= global microsatellite unstable MANTIS>0.4, MMR.D = loss of MMR protein expression, MutL = loss of expression of MLH1 and/or PMS2, MutS = loss of expression of MSH2 and/or MSH6.

**Supplementary Table 1.** **Demographics and clinical information of patients included in the study.**

LS = Lynch syndrome, LLS = Lynch-like syndrome, BRAF V600E = tumors with point mutation, M = male, F = female, WEA = White European Americans, AA = African Americans, SE = Spanish Europeans, CRC =colorectal cancer, FDRCRC = first degree relative with CRC, FDR/SDR LS tumor = first or second degree relative with a LS-related tumor, L = CRC left-sided, R = CRC right-sided.

**Supplementary Table 2. Sample origin and type of sample**

CCC = Chicago Colorectal Cancer Consortium, DFBI = Dana Farber Cancer Institute , EPI = EPICOLON Consortium, HGUA = Hospital General Universitario de Alicante in Spain, UMICH = University of Michigan, Yale = Yale New Haven Hospital, FFT = fresh-frozen tissue, FFPE = formalin-fixed paraffin embedded.

**Supplementary Table 3: Clinical and molecular information for the independent set of dMMR tumors.**

A) Clinical and molecular information: LLS = Lynch-like syndrome, MSI.BRAF = Microsatellite instable with BRAF V600E, IHC= loss of MMR protein expression , CRC= colorectal cancer, L = CRC left-sided, R = CRC right-sided, Age Dx = age of diagnosis, g-MSI = globally microsatellite instable by MANTIS, g-MSS = globally microsatellite stable by MANTIS, FDR.CRC = first degree relative with CRC, FDR/SDR LS tumor = first or second degree relative with a LS-related tumor. B) Signature contribution

**Supplementary Table 4. Sequencing metrics.**

**Supplementary Table 5. Comparison of clinical and molecular features of LS and LLS tumors between clusters**

LLS = Lynch-like syndrome, LS = Lynch syndrome, MSI.BRAF = Microsatellite instable with BRAF V600E, VUS = variant of uncertain significance, MutL = loss of expression of MLH1 and/or PMS2, MutS = loss of expression of MSH2 and/or MSH6, M = male, F = female, CRC = colorectal, L = CRC left-sided, R = CRC right-sided, FH = family history, ≥ 1FDR = One or more first degree relative, Sig = signature, TMB=Tumor Mutational Burden (total mutations/Mb).

ns = FDR >0.05.

**Supplementary Table 6. Patients with tumors with global exome microsatellite stability**

MSI PCR= MSI-H status by PCR, IHC= loss of MMR protein expression, Germline MMR= germline mutation in an MMR gene, Somatic MMR= somatic mutation in an MMR gene, Inact= MMR gene biallelic inactivation, Tumor location (age)= R right-side colon, L left-side colon, age of diagnosis, FDR and SDR LS-related tumor= first or second degree relative with a LS-related tumor/ total number of family members , CRC = colorectal cancer, OV = ovarian cancer, STOM = stomach cancer, BLA = bladder cancer, TMB=Tumor Mutational Burden (total mutations/Mb).

**Supplementary Table 7. Mutational events (SNV and LOH) in colorectal cancer driver genes by cluster.**

**Supplementary Table 8. Most recurrent HLA-I-A,B,C alleles among patients with dMMR tumors**

WEA = White European Americans, AA = African Americans, SE = Spanish Europeans.

**Supplementary Table 1.** **Demographics and clinical information of patients included in the study**

| **Phenotype** | **Age of Diagnosis** | **Sex** | | **Ethnicity** | | | **Family History** | | | **Tumor Location** | |
| --- | --- | --- | --- | --- | --- | --- | --- | --- | --- | --- | --- |
|  | Median  ± SD | M | F | WEA | AA | SE | None | FDR CRC | FDR/SDR LS tumor | R | L |
| LS  *(N= 25)* | 62 ± 13 | 60% (15/25) | 40% (10/25) | 36% (9/25) | 12% (3/25) | 52% (13/25) | 20% (4/20) | 35% (7/20) | 80% (16/20) | 58.3% (14/24) | 41.6% (10/24) |
| LLS  *(N= 35)* | 54 ± 13 | 57% (20/35) | 43% (15/35) | 28.6% (10/35) | 14.3% (5/35) | 57.1% (20/35) | 34.4% (11/32) | 40.6% (13/32) | 65.5% (21/32) | 76.6% (23/30) | 23.3% (7/30) |
| BRAF V600E  *(N= 43)* | 79 ± 9 | 28% (12/43) | 72% (31/43) | 87% (27/31) | 10% (3/31) | 3% (1/31) | 78% (29/37) | 22% (8/37) | NA | 98% (41/42) | 2% (1/42) |

**Supplementary Table 2. Sample origin and type of sample**

| **ID** | **Center** | **Sample type** | **ID** | **Center** | **Sample type** |
| --- | --- | --- | --- | --- | --- |
| PC1101 | CCC | FFT | 13112 | HGUA | FFPE |
| PC831 | CCC | FFT | 13113 | HGUA | FFPE |
| PC871 | CCC | FFT | 13114 | HGUA | FFPE |
| PC971 | CCC | FFT | 13115 | HGUA | FFPE |
| PS33 | CCC | FFT | 13116 | HGUA | FFPE |
| PS69 | CCC | FFT | 13120 | HGUA | FFPE |
| PV19 | CCC | FFT | 13122 | HGUA | FFPE |
| PV5 | CCC | FFT | 13123 | HGUA | FFPE |
| PV74 | CCC | FFT | 13124 | HGUA | FFPE |
| U27 | CCC | FFT | CL02138 | HGUA | FFPE |
| U43 | CCC | FFT | EL105563 | HGUA | FFPE |
| 23479H | DFCI | FFPE | EL56386 | HGUA | FFPE |
| 40041E | DFCI | FFPE | v860905 | HGUA | FFPE |
| A51080 | DFCI | FFPE | v860909 | HGUA | FFPE |
| J50801 | DFCI | FFPE | v860914 | HGUA | FFPE |
| K47750 | DFCI | FFPE | 5643 | UMICH | FFPE |
| LF61423 | DFCI | FFPE | 11032 | UMICH | FFPE |
| M38766 | DFCI | FFPE | 397 | Yale | FFPE |
| N25934 | DFCI | FFPE | 441 | Yale | FFPE |
| S047659 | DFCI | FFPE | S10-29135 | Yale | FFPE |
| 469 | EPI | FFPE | S10-4557T1.3ascd | Yale | FFPE |
| 4065 | EPI | FFPE | S11-10596s5 | Yale | FFPE |
| 5397 | EPI | FFPE | S11-25003 | Yale | FFPE |
| 5401 | EPI | FFPE | S11-32114 | Yale | FFPE |
| 6141 | EPI | FFPE | S12-15533 | Yale | FFPE |
| 6576 | EPI | FFPE | S12-7578 | Yale | FFPE |
| 6578 | EPI | FFPE | S13-24148 | Yale | FFPE |
| 6836 | EPI | FFPE | S13-25384 | Yale | FFPE |
| 7506 | EPI | FFPE | S13-28104 | Yale | FFPE |
| 7580 | EPI | FFPE | S14-25986 | Yale | FFPE |
| 15034 | EPI | FFPE | S14-26004 | Yale | FFPE |
| 13124Sp | EPI | FFPE | S15-13990 | Yale | FFPE |
| 83EPI2 | EPI | FFPE | S16-32637 | Yale | FFPE |
| 13102 | HGUA | FFPE | SR14-10903 | Yale | FFPE |
| 13105 | HGUA | FFPE | SR14-12774 | Yale | FFPE |
| 13107 | HGUA | FFPE | SR17-6975 | Yale | FFPE |
| 13111 | HGUA | FFPE | SR19-11388 | Yale | FFPE |

**Supplementary Table 3: Clinical and molecular information for the independent set of dMMR tumors.**

A. Clinical and tumor information

| **ID** | **Phenotype** | **IHC** | **CRC Location** | **Age Dx** | **Sex** | **Somatic hit** | **MANTIS Status** | **FDR CRC** | **FDR/SDR LS tumor** | **Tumor purity** |
| --- | --- | --- | --- | --- | --- | --- | --- | --- | --- | --- |
| 88172896 | LLS | MSH2 MSH6 | R | 47 | F | NO | g-MSI | NO | YES | 28.9% |
| 88180059 | LLS | PMS2 | L | 45 | M | MLH1 LOH | g-MSS | YES | NO | 35.7% |
| 88181153 | MSI.BRAF | MLH1  PMS2 | NA | NA | NA | MSH6  c.2864dupC p.T955fs | g-MSI | NA | NA | 36.0% |
| 88192310 | LLS | MLH1  PMS2 | L | 49 | M | MLH1 LOH and c.G193A:p.G65S | g-MSI | YES | YES | 33.3% |
| 88200138 | LLS | MLH1  PMS2 | R | 74 | M | NO | g-MSI | YES | YES | 29.80% |
| 88200284 | LLS | MSH2 MSH6 | L | 33 | M | MSH2  c.1386+3_1386+4insGTC | g-MSI | NO | YES | 44.2% |

B. Signature contribution

| **Sample** | **SBS1** | **SBS6** | **SBS14** | **SBS15** | **SBS20** | **SBS21** | **SBS26** | **SBS44** |
| --- | --- | --- | --- | --- | --- | --- | --- | --- |
| 88172896 | 630.12 | 297.43 | 10.61 | 626.88 | 33.20 | 0.00 | 316.76 | 0.00 |
| 88180059 | 137.37 | 11.22 | 0.00 | 0.00 | 0.00 | 0.00 | 68.29 | 0.00 |
| 88181153 | 233.90 | 281.59 | 62.05 | 1436.6 | 42.33 | 0.00 | 340.39 | 0.00 |
| 88192310 | 98.44 | 120.53 | 0.00 | 457.21 | 0.64 | 0.00 | 231.04 | 0.00 |
| 88200138 | 391.25 | 0.00 | 11.08 | 600.78 | 122.27 | 0.00 | 126.14 | 0.00 |
| 88200284 | 362.58 | 326.82 | 42.24 | 725.98 | 145.08 | 45.58 | 113.23 | 0.00 |

**Supplementary Table 4: Sequencing metrics**

| **Name** | **Tumor Purity** | **Mean Coverage** | **Name** | **Tumor Purity** | **Mean Coverage** | **Name** | **Tumor Purity** | **Mean Coverage** |
| --- | --- | --- | --- | --- | --- | --- | --- | --- |
| 397 | 80.9% | 196.1 | 13124 | 50.2% | 89.9 | S11-32114 | 70.3% | 107.3 |
| 441 | 81.6% | 199.7 | 15034 | 60.7% | 31.2 | S12-15533 | 27.1% | 188.9 |
| 469 | 37.5% | 112.2 | 13124Sp | 59% | 225.8 | S12-7578 | 59.6% | 190.4 |
| 4065 | 98.0% | 11.6 | 23479H | 43.2% | 61.2 | S13-24148 | 48.5% | 160.5 |
| 5397 | 67.0% | 85 | 40041E | 59.2% | 96.4 | S13-25384 | 67.40% | 118.9 |
| 5401 | 49.6% | 76.9 | EL56386 | 43.8% | 68 | S13-28104 | 44.6% | 102.2 |
| 5643 | 40.7% | 97.6 | J50801 | 39.9% | 131.3 | S14-25986 | 65.6% | 214 |
| 6141 | 56% | 48.9 | K47750 | 76.6% | 90.9 | S14-25986.211 | 64.4% | 117.1 |
| 6576 | 54.6% | 103.7 | LF61423 | 42.1% | 78 | S14-26004 | 68.1% | 128 |
| 6578 | 51.2% | 151.3 | M38766 | 45.0% | 105 | TU27 | 69% | 163.3 |
| 6836 | 64% | 109.6 | N25934 | 73.9% | 83.9 | S16-32637 | 61.8% | 88.2 |
| 7506 | 48.7% | 67.9 | PC1101 | 49.5% | 183.3 | SR14-10903 | 35.7% | 101.4 |
| 7580 | 40.5% | 118.9 | PC831 | 47.9% | 155.4 | SR14-12774 | 52.4% | 143.8 |
| 11032 | 48.3% | 146.4 | PC871 | 73.2% | 169.2 | SR17-6975 | 75.1% | 127.4 |
| 13102 | 46.5% | 103.6 | PC971 | 44% | 237.6 | SR19-11388 | 56.4% | 243.9 |
| 13105 | 57.1% | 49.9 | PS33 | 54.8% | 228.4 | EL105563 | 76.3% | 116.2 |
| 13107 | 59.9% | 99.9 | PS69 | 44.2% | 128.4 | S15-13990 | 43.4% | 83.2 |
| 13111 | 80.4% | 94.8 | PV19 | 65.0% | 149.7 | 83EPI | 54% | 143.5 |
| 13112 | 76.8% | 120.1 | PV5 | 66.9% | 140 | A51080 | 72.8% | 188.9 |
| 13113 | 39.5% | 58.4 | PV74 | 59.2% | 144.1 | CL02138 | 45.5% | 67.8 |
| 13114 | 64.4% | 82.6 | S047659 | 49.5% | 39.3 | EL01021 | 55.9% | 101.8 |
| 13115 | 50.0% | 64.4 | S10-29135 | 70.2% | 153.1 | TU43 | 75.8% | 177.9 |
| 13116 | 40.0% | 97.7 | S10-4557.3 | 60.2% | 189.4 | v860905 | 51.3% | 79.7 |
| 13120 | 61.2% | 105.7 | S11-10596.5 | 68.1% | 128 | v860909 | 41.8% | 85.6 |
| 13122 | 40.9% | 157.8 | S11-10596.9 | 52.5% | 149.2 | v860914 | 82.1% | 85.3 |
| 13123 | 38.7% | 131.8 | S11-25003 | 52.8% | 140.2 | **Median ± sd** | **54.7 ± 0.14** | **105.7 ± 50.2** |

.

**Supplementary Table 5. Comparison of clinical and molecular features of LS and LLS tumors between clusters**

|  |  | **Cluster1** | | **Cluster2** | | **Cluster3** | | **FDR** |
| --- | --- | --- | --- | --- | --- | --- | --- | --- |
|  |  | n=31 | | n=23 | | n=6 | |  |
| **Phenotype** | LLS | 61% | (19) | 52% | (12) | 67% | (4) | ns |
|  | LS | 39% | (12) | 48% | (11) | 33% | (2) |  |
| **Loss Expression MMR*** | MutL | 65% | (20) | 27% | (6) | 67% | (4) | 5.5x10^-5^ |
|  | MutS | 35% | (11) | 73% | (16) | 33% | (2) |  |
| **Sex** | M | 65% | (20) | 52% | (12) | 50% | (3) | ns |
|  | F | 35% | (11) | 48% | (11) | 50% | (3) |  |
| **Age** |  | 55±13 | | 50±13 | | 74±9 | | ns |
| **(median ±sd)** |  |  |  |  |  |  |  |  |
| **Location** | L | 34% | (10) | 39% | (7) | 17% | (1) | ns |
|  | R | 66% | (19) | 61% | (11) | 83% | (5) |  |
| **Stage** | I/II | 45% | (14) | 43% | (10) | 100% | (6) | ns |
|  | III/IV | 26% | (8) | 17% | (4) | 0% | (0) |  |
|  | NA | 29% | (9) | 4% | (9) | 0% | (0) |  |
| **FH of CRC** | none | 66% | (19) | 58% | (11) | 67% | (4) | ns |
|  | ≥ 1FDR | 44% | (10) | 42% | (8) | 33% | (2) | ns |
| **Signature** | SBS1 | 226±235 | | 507±315 | | 193±83 | | 0.026 |
| **Composition (%)** | SBS6 | 180±125 | | 127±111 | | 0±25 | | 0.003 |
|  | SBS14 | 4±35 | | 10±32 | | 0±8 | | ns |
|  | SBS15 | 295±267 | | 414±412 | | 751±383 | | 0.049 |
|  | SBS20 | 57±61 | | 47±83 | | 0±28 | | 0.026 |
|  | SBS21 | 0±161 | | 0±10 | | 0±30 | | ns |
|  | SBS26 | 85±153 | | 57±71 | | 109±137 | | ns |
|  | SBS44 | 0±0 | | 0±0 | | 0±0 | | NA |
| **Tumor purity (%)** | 53±13 | | | 50±16 | | 60±13 | | ns |
| **# events** | TMB | 40±22 | | 45±38 | | 38±13 | | ns |
| **(median ±sd)** | frameshifts | 78±115 | | 68±100 | | 48±97 | | ns |
| **MANTIS score** | 0.48±0.13 | | | 0.48±0.15 | | 0.49±0.06 | | ns |

*one tumor in Cluster 2 did not show loss of MMR expression, but was MSI-H

**Supplementary Table 6: Patients with tumors with global microsatellite stability**

| **ID** | **MSI PCR** | **IHC** | **Germline MMR** | **Somatic MMR** | **Biallelic** | **Tumor Location (age)** | **FDR and SDR LS-related tumor** | | **TMB** | **#fs** |
| --- | --- | --- | --- | --- | --- | --- | --- | --- | --- | --- |
| 13107 | MSI | MSH6 | c.778delG:p.D260fs MSH6 | c.C1093T:p.R365X  MSH6 | yes | L (68) | 2/13 | CRC (75) | 44.3 | 6 |
| 13113 | MSI | PMS2 | c.C1882T.p.R628X  PMS2 | no mutation |  | R (62) | 2/28 | OV (60) | 30.4 | 2 |
| PC971 | MSI | MSH6 | c.1312_1313del:p.F438fs MSH6 | c.C2374T:p.R792X  MSH6 | yes | L (78) | NA | NA | 75.4 | 16 |
| 13116 | NA | MSH2  MSH6 | c.C1552T:p.Q518X  MSH2 | c.2459-2A>G  MSH2 | yes | R (59) | NA | NA | 31.4 | 1 |
| 13122 | NA | MSH2  MSH6 | c.C1216T:p.R406X  MSH2 | no mutation |  | L (65) | 3/21 | STOM (NA,NA) | 40.9 | 28 |
| 7580 | MSI | MLH1  PMS2 | no mutation | MLH1 LOH and c.T1520A:p.L507X | yes | L (56) | 0 |  | 36.0 | 3 |
| 5397 | MSI | MSH6 | no mutation | MSH2  p.K871N |  | R (47) | 4/8 | CRC, STOM, BLA (73, 75, 83, NA) | 193.4 | 21 |
| M38766 | MSI | MSH2 MSH6 | no mutation | no mutation |  | R (58) | 5/10 | CRC (60, 70,62, 60, 68) | 13.9 | 0 |
| 5401 | MSI | MSH6 | no mutation | LOH  MLH1, MSH6, MSH2 |  | L (56) | 2/7 | CRC, STOM (74, 85) | 8.7 | 2 |

**Supplementary Table 7: Mutational events (SNV and LOH) in colorectal cancer driver genes by cluster.**

|  | **Cluster 1** | | **Cluster 2** | | **Cluster 3** | |
| --- | --- | --- | --- | --- | --- | --- |
| **Gene** | **Events** | **Frequency (%)** | **Events** | **Frequency (%)** | **Events** | **F Frequency (%)** |
| *ACVR2A* | 11 | 13.4 | 5 | 10.9 | 1 | 5.6 |
| *AMER1* | 6 | 7.3 | 7 | 15.2 | 3 | 16.7 |
| *APC* | 26 | 31.7 | 9 | 19.6 | 5 | 27.8 |
| *ARID1A* | 21 | 25.6 | 6 | 13.0 | 7 | 38.9 |
| *BRAF* | 26 | 31.7 | 6 | 13.0 | 3 | 16.7 |
| *CTNNB1* | 28 | 34.1 | 8 | 17.4 | 2 | 11.1 |
| *FBXW7* | 24 | 29.3 | 9 | 19.6 | 3 | 16.7 |
| *GNAS* | 19 | 23.2 | 6 | 13.0 | 3 | 16.7 |
| *KRAS* | 22 | 26.8 | 9 | 19.6 | 5 | 27.8 |
| *NRAS* | 16 | 19.5 | 1 | 2.2 | 2 | 11.1 |
| *PCBP1* | 15 | 18.3 | 3 | 6.5 | 1 | 5.6 |
| *PIK3CA* | 21 | 25.6 | 6 | 13.0 | 3 | 16.7 |
| *PTEN* | 18 | 22.0 | 6 | 13.0 | 1 | 5.6 |
| *SMAD2* | 15 | 18.3 | 6 | 13.0 | 1 | 5.6 |
| *SMAD4* | 18 | 22.0 | 4 | 8.7 | 0 | 0.0 |
| *SOX9* | 23 | 28.0 | 5 | 10.9 | 3 | 16.7 |
| *TCF7L2* | 21 | 25.6 | 5 | 10.9 | 1 | 5.6 |
| *TGIF1* | 15 | 18.3 | 1 | 2.2 | 0 | 0.0 |
| *TP53* | 21 | 25.6 | 5 | 10.9 | 2 | 11.1 |
| *ZFP36L2* | 20 | 24.4 | 3 | 6.5 | 2 | 11.1 |

**Supplementary Table 8: Most recurrent HLA-I-A,B,C alleles among patients with dMMR tumors**

A. Cluster 1

| **HLA-A01:01** | **HLA-A02:01** | **HLA-B44:02** | **HLA-B07:02** | **HLA-C05:01** | **HLA-C07:02** | **ID** | **Ethnicity** |
| --- | --- | --- | --- | --- | --- | --- | --- |
| 0/0 | 0/0 | 0/1 | 0/1 | 0/0 | 0/1 | 397 | NA |
| 0/0 | 0/0 | 0/0 | 0/0 | 0/0 | 0/0 | 6576 | SE |
| 0/0 | 0/0 | 0/0 | 1/1 | 0/0 | 0/0 | 6578 | SE |
| 0/1 | 0/0 | 0/0 | 0/0 | 0/1 | 0/0 | 7506 | SE |
| 0/0 | 0/1 | 0/1 | 0/0 | 0/1 | 0/0 | 7580 | SE |
| 0/0 | 0/0 | 0/0 | 0/0 | 0/1 | 0/0 | 11032 | WEA |
| 0/0 | 0/0 | 0/0 | 0/0 | 0/0 | 0/1 | 13102 | SE |
| 0/0 | 0/0 | 0/0 | 0/0 | 0/0 | 0/0 | 13105 | SE |
| 0/1 | 0/0 | 0/0 | 0/0 | 0/0 | 1/1 | 13107 | SE |
| 0/0 | 0/0 | 0/0 | 0/0 | 0/0 | 0/0 | 13112 | SE |
| 0/0 | 0/0 | 0/0 | 0/0 | 0/0 | 0/0 | 13123 | SE |
| 0/1 | 0/0 | 0/0 | 0/0 | 0/0 | 0/0 | 40041E | AA |
| 0/0 | 0/0 | 0/0 | 0/0 | 0/1 | 0/0 | 83EPI2 | SE |
| 0/0 | 0/0 | 0/0 | 0/1 | 0/0 | 1/1 | A51080 | WEA |
| 0/0 | 0/0 | 0/0 | 0/0 | 0/0 | 0/0 | EL105563 | SE |
| 0/0 | 0/1 | 0/0 | 1/1 | 0/0 | 0/1 | K47750 | WEA |
| 0/0 | 0/0 | 0/0 | 0/0 | 0/0 | 0/0 | LF61423 | SE |
| 0/0 | 0/1 | 0/0 | 0/0 | 0/0 | 0/0 | M38766 | WEA |
| 0/0 | 0/1 | 1/1 | 0/0 | 0/0 | 0/0 | N25934 | WEA |
| 0/1 | 0/1 | 0/1 | 0/0 | 0/0 | 0/0 | PC1101 | AA |
| 0/0 | 0/0 | 0/0 | 0/0 | 0/0 | 0/1 | PC831 | WEA |
| 0/0 | 0/0 | 0/1 | 0/0 | 0/0 | 0/1 | PC871 | AA |
| 0/0 | 0/1 | 0/0 | 1/1 | 0/1 | 0/0 | PS33 | WEA |
| 0/0 | 1/1 | 0/0 | 0/0 | 0/0 | 0/0 | PV19 | AA |
| 0/0 | 1/1 | 0/0 | 0/0 | 0/0 | 0/0 | PV5 | WEA |
| 0/0 | 0/0 | 0/1 | 0/0 | 0/0 | 0/0 | S047659 | WEA |
| 0/0 | 0/1 | 0/0 | 0/0 | 0/0 | 0/1 | S10-4557 | WEA |
| 0/0 | 1/1 | 0/1 | 0/0 | 0/0 | 0/0 | S11-10596.15 | other |
| 0/0 | 0/1 | 0/0 | 0/0 | 0/0 | 0/0 | S11-10596.19 | other |
| 0/0 | 1/1 | 0/0 | 0/0 | 0/0 | 0/1 | S11-25003 | WEA |
| 0/0 | 0/1 | 0/0 | 0/1 | 0/0 | 0/0 | S12-15533 | WEA |
| 0/1 | 0/0 | 0/0 | 0/0 | 0/0 | 0/1 | S12-7578 | WEA |
| 0/1 | 0/0 | 0/1 | 0/0 | 0/1 | 0/0 | S13-24148 | WEA |
| 0/1 | 0/1 | 0/0 | 1/1 | 0/0 | 0/0 | S13-25384 | WEA |
| 0/0 | 0/0 | 0/0 | 0/0 | 0/0 | 0/0 | S13-28104 | WEA |
| 0/0 | 0/0 | 0/0 | 0/0 | 0/0 | 0/0 | S14-25986 | WEA |
| 0/0 | 0/0 | 0/0 | 0/0 | 0/0 | 0/0 | S14-26004 | WEA |
| 0/0 | 0/0 | 0/0 | 0/0 | 0/0 | 0/0 | S15-13990 | WEA |
| 0/0 | 0/0 | 0/0 | 0/1 | 0/1 | 0/0 | SR14-10903 | WEA |
| 0/0 | 1/1 | 0/0 | 0/0 | 0/0 | 0/1 | SR14-12274 | WEA |
| 0/0 | 0/1 | 0/0 | 0/0 | 0/1 | 0/0 | SR19-11388 | WEA |
| 0/0 | 0/1 | 0/0 | 0/0 | 0/0 | 0/1 | TU27 | AA |
| 0/1 | 0/0 | 0/0 | 0/0 | 0/0 | 0/0 | U43 | AA |
| 0/1 | 0/0 | 0/0 | 0/1 | 0/0 | 0/0 | v860905 | SE |

B. Cluster 2

| **HLA-A01:01** | **HLA-A02:01** | **HLA-B44:02** | **HLA-B07:02** | **HLA-C05:01** | **HLA-C07:02** | **ID** | **Ethnicity** |
| --- | --- | --- | --- | --- | --- | --- | --- |
| 0/0 | 0/0 | 0/0 | 0/0 | 0/0 | 0/0 | 441 | AA |
| 0/0 | 0/0 | 0/0 | 0/0 | 0/0 | 0/0 | 5397 | SE |
| 0/1 | 0/0 | 0/1 | 0/0 | 0/1 | 0/0 | 5401 | SE |
| 0/0 | 0/0 | 0/0 | 0/0 | 0/0 | 0/0 | 6141 | SE |
| 0/0 | 0/1 | 0/0 | 0/0 | 0/1 | 0/0 | 13111 | SE |
| 0/1 | 0/0 | 1/1 | 0/0 | 0/1 | 0/0 | 13113 | SE |
| 0/1 | 0/0 | 0/1 | 0/0 | 0/1 | 0/0 | 13114 | SE |
| 0/0 | 0/0 | 0/0 | 0/0 | 0/0 | 0/0 | 13115 | SE |
| 0/0 | 0/1 | 0/1 | 0/0 | 0/1 | 0/0 | 13116 | SE |
| 0/1 | 0/1 | 0/0 | 1/1 | 0/1 | 0/1 | 13120 | SE |
| 1/1 | 0/0 | 0/0 | 0/0 | 0/0 | 0/0 | 13122 | SE |
| 0/0 | 0/0 | 0/1 | 0/0 | 0/0 | 0/0 | 13124 | WEA |
| 1/1 | 0/0 | 0/1 | 0/0 | 0/1 | 0/0 | 860909 | SE |
| 0/1 | 0/0 | 0/0 | 0/0 | 0/0 | 0/0 | 860914 | SE |
| 0/1 | 0/0 | 0/0 | 0/1 | 0/0 | 0/0 | 23470H | WEA |
| 0/0 | 0/0 | 0/0 | 0/0 | 0/1 | 0/0 | CL02138 | SE |
| 0/0 | 0/0 | 0/0 | 0/0 | 0/0 | 0/0 | EL56386 | SE |
| 0/0 | 0/0 | 0/0 | 0/0 | 0/0 | 0/0 | PC971 | AA |
| 0/0 | 0/1 | 0/0 | 0/0 | 0/0 | 0/0 | PS69 | AA |
| 0/0 | 0/0 | 0/0 | 0/0 | 0/0 | 0/0 | PV74 | AA |
| 0/0 | 0/0 | 0/0 | 0/0 | 0/0 | 0/0 | S11-32114 | WEA |
| 1/1 | 0/0 | 0/1 | 0/0 | 0/1 | 0/0 | S16-32637 | NA |

C. Cluster 3

| **HLA-A01:01** | **HLA-A02:01** | **HLA-B44:02** | **HLA-B07:02** | **HLA-C05:01** | **HLA-C07:02** | **ID** | **Ethnicity** |
| --- | --- | --- | --- | --- | --- | --- | --- |
| 0/0 | 0/0 | 0/0 | 1/1 | 0/1 | 0/1 | 469 | SE |
| 0/0 | 0/1 | 0/0 | 0/0 | 0/0 | 0/0 | 5643 | NA |
| 0/0 | 0/1 | 0/0 | 0/0 | 0/1 | 0/0 | 6836 | SE |
| 0/1 | 0/0 | 0/0 | 0/0 | 0/0 | 0/0 | 15034 | SE |
| 1/1 | 0/0 | 0/0 | 0/0 | 0/0 | 0/0 | 13124Sp | SE |
| 0/0 | 0/1 | 0/0 | 0/0 | 0/0 | 0/0 | J50801 | WEA |
| 1/1 | 0/0 | 0/0 | 0/0 | 0/0 | 0/0 | S10-29315 | NA |
| 0/1 | 0/0 | 0/0 | 0/0 | 0/0 | 0/0 | S14.25986s211 | WEA |
| 0/1 | 0/1 | 0/0 | 0/1 | 0/1 | 0/1 | SR17-6975 | WEA |
