## Supplementary figures and images for "Mutational signature profiling classifies subtypes of clinically different mismatch repair deficient tumors with a differential immunogenic response potential"

### Supplementary Figure 1.tiff

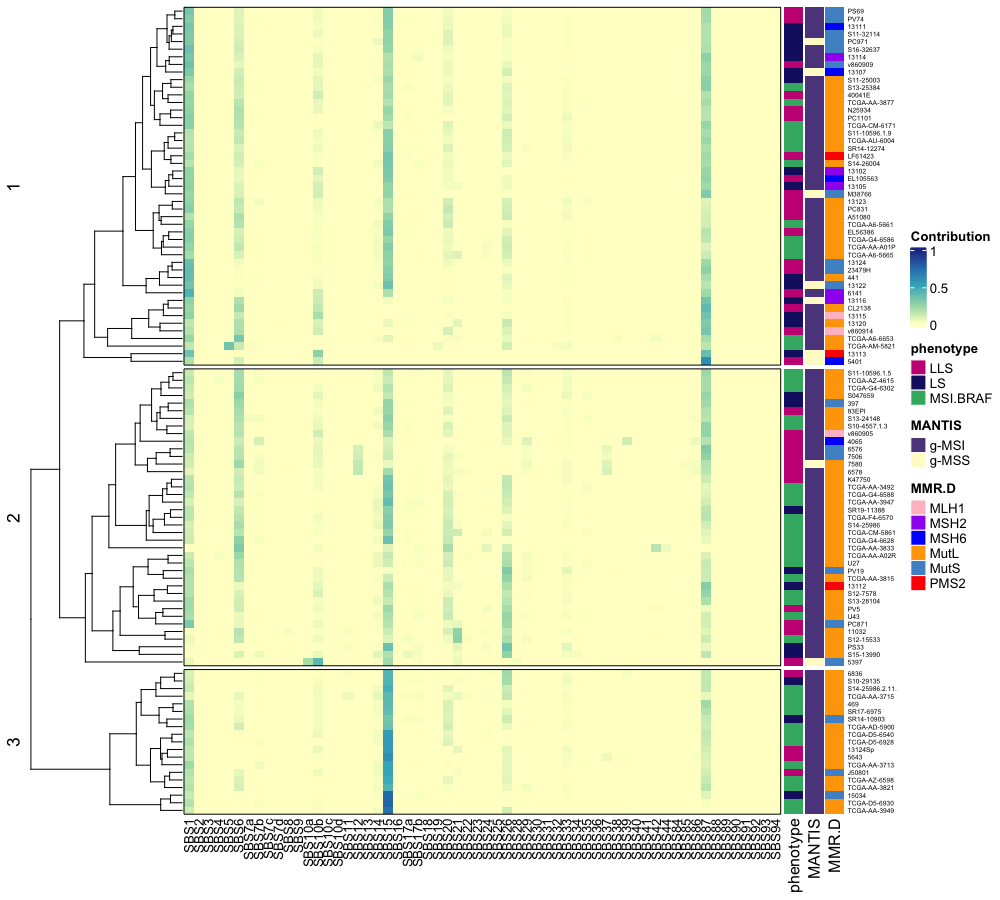

### Supplementary Figure 3

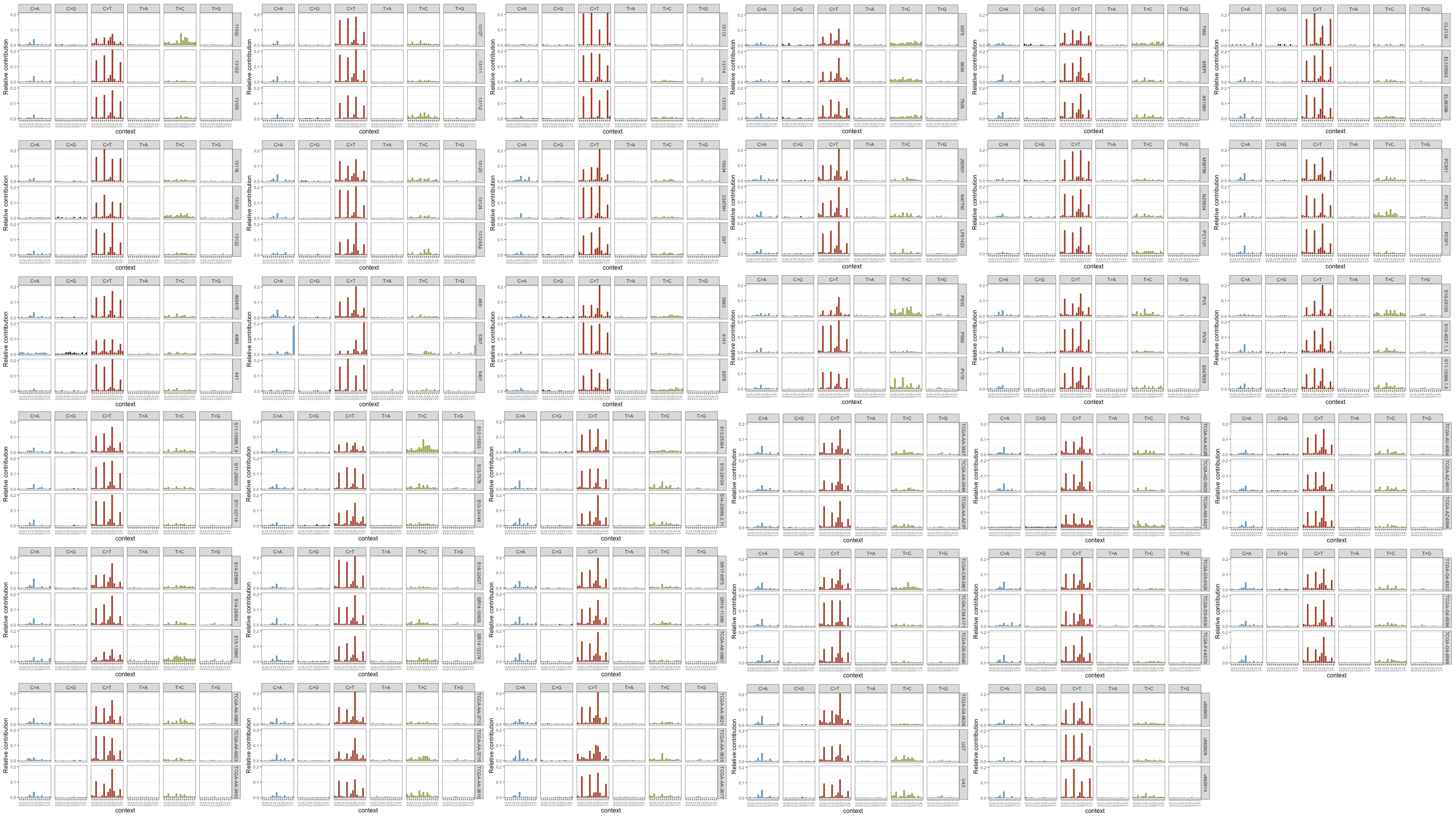

### Supplementary Figure 4.tiff

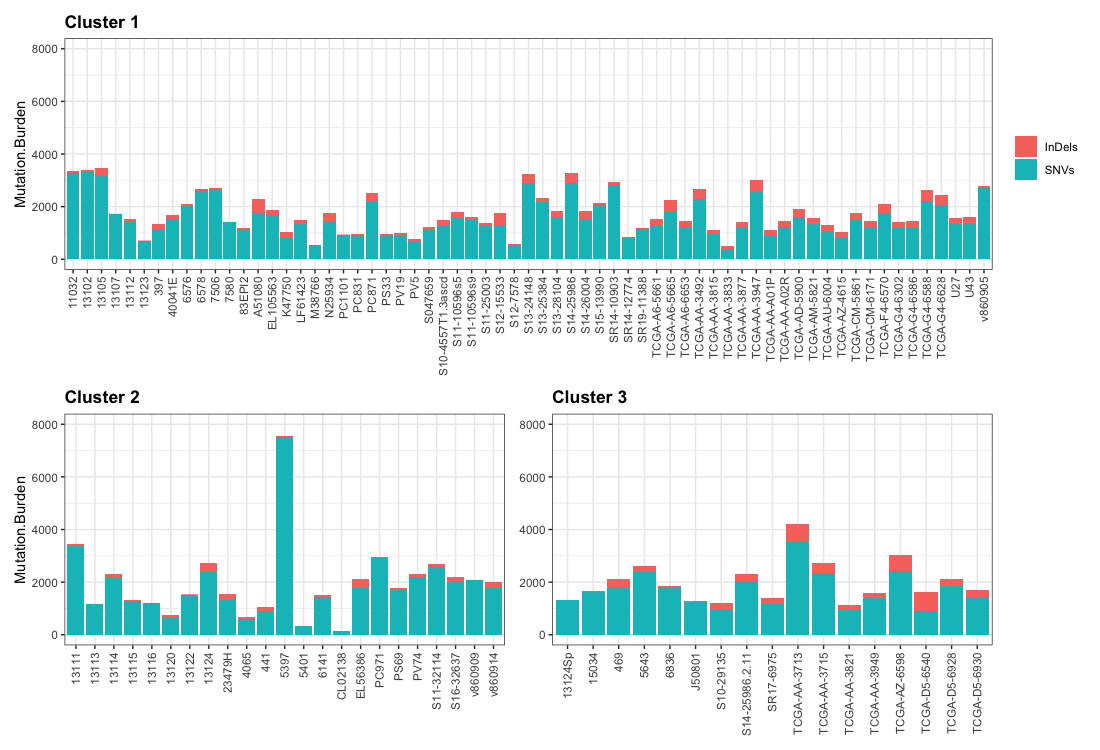

### Supplementary Figure 5 .tiff

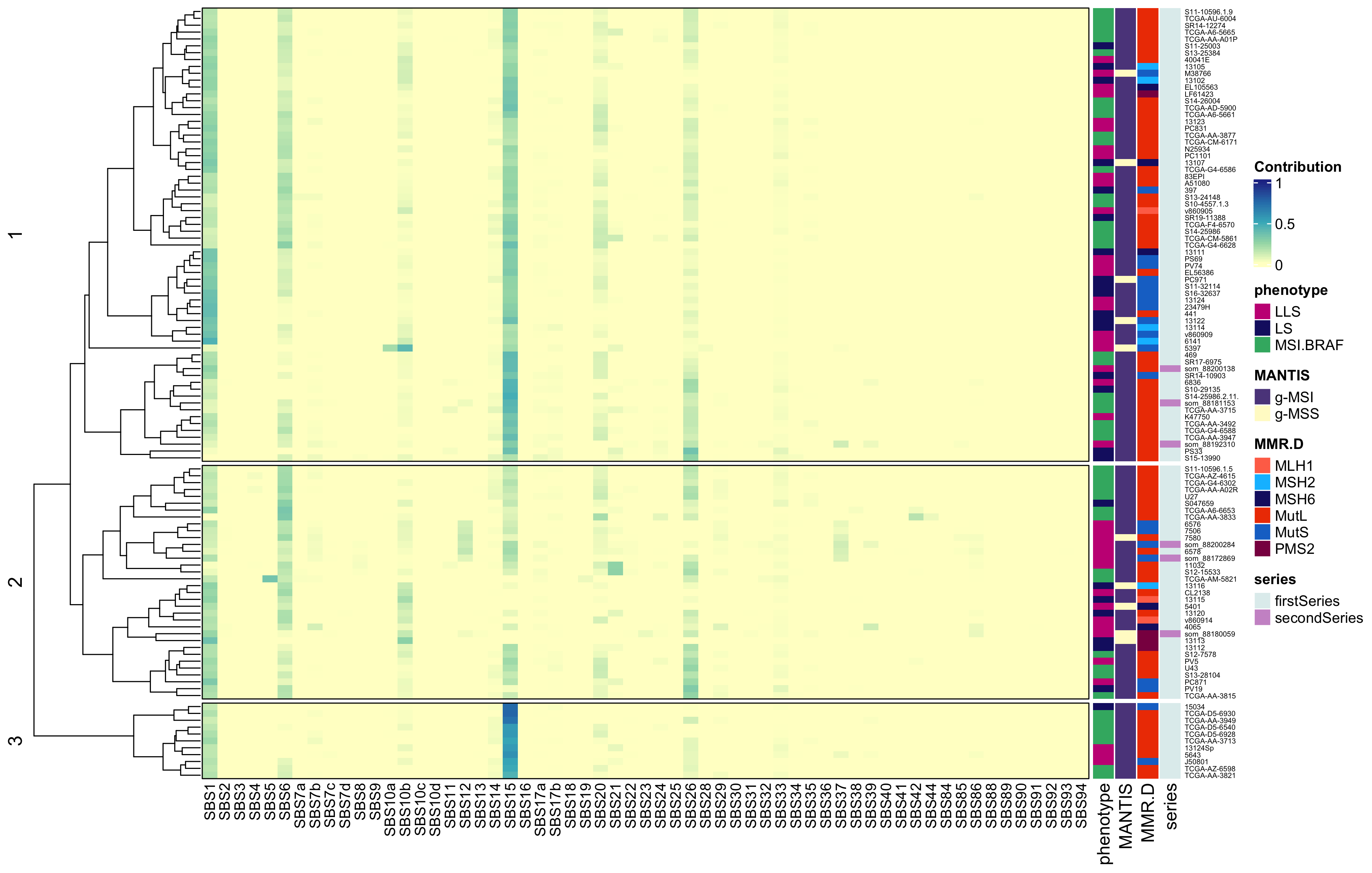
